# Developmental programming of adrenal chromaffin cell connexin plasticity by neonatal maternal separation

**DOI:** 10.64898/2026.06.16.732707

**Authors:** Pedro Segura-Chama, Vito S. Hernández, Limei Zhang

## Abstract

Adrenal chromaffin cells are key effectors of the sympathoadrenal response and play a central role in the organism’s adaptation to environmental and physiological challenges. While cholinergic and pituitary adenylate cyclase-activating polypeptide (PACAP)-dependent mechanisms have long been recognized as major regulators of catecholamine secretion, increasing evidence indicates that connexin-mediated gap junctional communication provides an additional and highly dynamic level of control. Whether early-life experience modifies the adult capacity of chromaffin-cell networks to undergo stress-induced connexin remodeling remains unclear.

Here, we examined adrenal medullary connexin expression in adult rats exposed to neonatal maternal separation (MS; 3 h daily, postnatal days 2-15) and later challenged with an 8-day unpredictable mild stress (UMS) protocol. Under basal adult conditions, MS did not produce an overt change in adrenal medullary Cx36 or Cx43 immunoreactivity relative to animal-facility-reared controls. In contrast, UMS increased connexin immunoreactivity in the adrenal medulla, and this response was amplified in animals with a history of MS. MS+UMS animals also displayed enhanced corticosterone responses to acute restraint stress. These findings suggest that neonatal MS does not impose a constitutively altered adult chromaffin-cell phenotype, but instead primes the future stress responsiveness of adrenal medullary connexin remodeling. We propose that chromaffin-cell gap junctions represent a substrate sensitive to stress history, through which developmental experience may influence sympathoadrenal and endocrine adaptation in adulthood.

## 1. Introduction

The sympathoadrenal system is a major effector of the acute stress response. Preganglionic splanchnic neurons activate adrenal chromaffin cells, which release catecholamines to support cardiovascular, metabolic, and behavioral adaptation during physiological challenge (Douglas 1968, Smith and Eiden 2012, Kanczkowski, Sue et al. 2016). This canonical view has traditionally emphasized synaptic control by acetylcholine (ACh) and PACAP. However, chromaffin cells are not isolated secretory units. Within the adrenal medulla, they are organized in compact clusters embedded among vascular sinusoids and splanchnic nerve terminals, creating a local architecture that favors coordinated activity (Perlman and Chalfie 1977, Iijima, Matsumoto et al. 1992, Kajiwara, Sand et al. 1997).

Gap junctional communication provides a mechanism through which chromaffin cells can operate as functional ensembles. Connexin-mediated coupling allows the spread of electrical and metabolic signals between neighboring cells and contributes to synchronized Ca2+ elevations and coordinated catecholamine release (Martin, Mathieu et al. 2001, Yamagami, Moritoyo et al. 2002, Colomer, Olivos Ore et al. 2008, Guérineau 2024). In this context, connexins act as conditional amplifiers: their contribution becomes most relevant when secretory demand is high, such as during intense splanchnic activity or repeated stress.

Chromaffin-cell coupling is developmentally and physiologically plastic. Neonatal chromaffin cells exhibit prominent local coupling before full maturation of splanchnic synaptic control, whereas adult coupling is dynamically regulated by synaptic activity, PACAP signaling, chronic stress, sex, aging, and cardiometabolic state (Martin, Mathieu et al. 2003, Colomer, Olivos-Ore et al. 2010, Hill, Lee et al. 2012, Desarmenien, Jourdan et al. 2013, Paille, Park et al. 2024). These observations raise the possibility that gap junctional communication may not only respond acutely to stress, but may also retain information about prior physiological or developmental history.

Early-life stress is a potent modifier of adult neuroendocrine function. In previous work using a mild maternal separation paradigm, we found persistent changes in the hypothalamic vasopressin system and related behavioral and neuroendocrine outcomes (Zhang, Hernandez et al. 2012, Hernandez, Hernandez et al. 2016, Zetter, Hernandez et al. 2021). Vasopressinergic signaling in the paraventricular nucleus can influence presympathetic neurons and thereby provides a plausible route by which early-life experience may alter later autonomic and adrenal recruitment (Son, Filosa et al. 2013). However, whether neonatal maternal separation modifies adult adrenal medullary connexin plasticity has not been directly tested.

Here, we asked whether neonatal maternal separation alters basal adult connexin expression in adrenal chromaffin cells or modifies the capacity of these cells to remodel connexin expression after a later stress challenge. We compared adult animal-facility-reared controls and maternally separated rats under basal conditions and after unpredictable mild stress (UMS). We further examined corticosterone responses to acute restraint stress as a functional endocrine readout of adrenal recruitment.

## 2. Materials and Methods

### 2.1 Animals and neonatal maternal separation procedure

Wistar rats bred in the local animal facility were used in this study. Animals were maintained under standard laboratory conditions, with a 12:12 h light-dark cycle, lights on at 07:00 h, controlled temperature of 22 ± 2 °C, and food and water available *ad libitum*. Adult female and male rats were paired for mating for two days. During the last week of gestation, pregnant females were single-housed in standard Plexiglas rat cages and left undisturbed until parturition.

All animal procedures were approved by the local bioethical and research committees of the School of Medicine of the National Autonomous University of México (CICUAL DI-FM-79/2020) and were conducted in accordance with institutional guidelines and with the principles described in the National Institutes of Health Guide for the Care and Use of Laboratory Animals (Committee 2011)

Neonatal maternal separation was performed using the same general procedure previously established in our laboratory for the study of hypothalamic vasopressin system regulation after maternal separation (Zhang, Hernández et al., 2012). On the day after parturition, designated postnatal day 2 (PND2), each litter was culled to 8 pups when possible. From PND2 to PND15, pups assigned to the maternal-separation group were separated from their dams for 3 h daily, between 09:00 and 12:00 h.

At the beginning of each separation session, pups were removed from the home cage by hand. To minimize olfactory disturbance, the experimenter’s hands were previously coated with fine bedding powder obtained from the corresponding home cage. Pups were then transferred to an adjacent room, placed individually in small boxes containing bedding, and maintained in a humidified incubator at 29 ± 1 °C for the duration of the 3 h separation period. After separation, pups were returned to their home cage and reunited with their respective dam.

Non-separated control litters, referred to here as animal-facility-reared controls, were left undisturbed except for routine cage maintenance and bedding changes twice per week. After weaning at PND 28, male and female offspring were assigned to basal or unpredictable mild stress (UMS) conditions. To minimize litter-related confounds, animals from multiple litters were distributed across experimental groups whenever possible. Animals were housed three per cage, under the above-mentioned standard conditions until testing in adulthood.

### 2.2 Adult unpredictable mild stress (UMS) protocol

At PND150, subsets of animal-facility-reared and maternally separated rats were exposed to an 8-day unpredictable mild stress protocol modified from (Zhang, Hernandez et al. 2008). The protocol consisted of alternating exposure to forced swim stress, sleep deprivation, restraint stress, and Morris water maze testing. Stressors were performed during the resting phase of the light–dark cycle, while the lights were on, and were applied at variable time points across days to reduce temporal predictability.

For forced swim stress, rats were placed individually for 5 min in a Plexiglas cylinder, 40 cm in diameter, filled with water maintained at 25 ± 1 °C to a depth that prevented escape and prevented the animal from touching the bottom with its tail or hindlimbs. After the session, animals were removed from the cylinder, gently dried, and returned to their home cage.

For sleep deprivation, rats were placed for 24 h in a water-filled chamber containing small elevated platforms. This configuration allowed animals to remain above the water while awake, whereas sleep-associated postural relaxation resulted in contact with the water and interruption of sleep. Water and room temperature were maintained at approximately 22 °C throughout the procedure. After the session, animals were returned to their home cages.

For Morris water maze exposure, rats were placed in a circular pool, 1.8 m in diameter, filled with water maintained at 22 ± 1 °C, and trained to locate a hidden escape platform, 15 cm in diameter. Each animal performed eight consecutive trials, with an intertrial interval of 10 min. The platform location was kept constant during the session, and animals were guided to the platform if they failed to locate it within one minute.

For restraint stress, rats were placed individually in restraining containers that limited major body movements without compressing the animal or interfering with respiration. Each restraint session lasted 30 min, after which animals were released and returned to their home cage.

The order of stressor exposure was varied across the 8-day protocol to minimize habituation and anticipation. Control animals assigned to the basal condition remained undisturbed in their home cages except for routine animal care.

### 2.3 Restraint stress and corticosterone measurement

For endocrine assessment, the last day of the unpredictable mild stress protocol consisted of restraint stress. Rats were placed individually in restraining containers that limited major body movements without compressing the animal or interfering with respiration. After 30 min of restraint, blood samples were obtained by tail-tip blood collection. Briefly, approximately 2 mm of the distal tail tip was clipped using sterile scissors or a sterile scalpel blade, and 500 µL of blood were collected into heparinized microcentrifuge tubes. Blood collection was completed as rapidly as possible to minimize additional handling-induced endocrine activation. Hemostasis was achieved by gentle pressure after sampling, and animals were monitored before being returned to their home cages.

Blood samples were centrifuged at 3000 × g for 10 min at 4 °C. Plasma was collected and stored at -20 °C until corticosterone measurement. Plasma corticosterone concentrations were determined using a commercial Corticosterone ELISA Kit (Abcam, Cat. No. AB323743), according to the manufacturer’s instructions. Rat plasma samples were diluted (1:4) in 1X Diluent S before assay. Standards and samples were assayed in duplicate. Absorbance was read at 450 nm using a microplate reader. Corticosterone concentrations were calculated from a standard curve and multiplied by the corresponding dilution factor.

According to the manufacturer, the minimum detectable corticosterone concentration was 0.33 ng/ml. The reported intra-assay and inter-assay coefficients of variation were 5.49% and 10.6%, respectively.

### 2.4 Tissue collection and adrenal processing

At PND100, rats were deeply anesthetized with pentobarbital 150 mg/kg and transcardially perfused with 0.9% saline, followed by fixative solution containing 4% formaldehyde in 0.1 M phosphate buffer (PB) and 15% v/v saturated picric acid solution, adjusted to pH 7.4. Adrenal glands were dissected, and rinsed in 0.1 M PB. Tissue was then embedded in gelatin and immediately sectioned at 70 µm using a Leica VT1000 vibratome. Free-floating adrenal sections were processed for immunohistochemical detection of connexins in the adrenal medulla.

### 2.5 Immunohistochemistry for Cx36 and Cx43

Free-floating adrenal sections were processed for immunofluorescent detection of connexin 36 (Cx36) and connexin 43 (Cx43). Cx36 and Cx43 immunostaining was performed in separate section series. Briefly, sections were rinsed in Tris-buffered saline (TBS) and incubated for 2 h at room temperature in blocking solution containing 10% normal donkey serum (NDS) in TBS with 0.3% Triton X-100 (TBST). Sections were then incubated for 48 hr. at 4 °C with rabbit anti-Cx36 (Invitrogen, Cat. 36-4600) or rabbit anti-Cx43 (Invitrogen, Cat. 71-0700) primary antibodies, diluted 1:200 in Tris-Buffered Saline with Triton 100x 0.3% (TBST) containing 1% normal donkey serum (NDS, Jackson ImmunoResearch, Cat. 017-000-121).

After primary antibody incubation, sections were washed three times for 10 min each in TBST and incubated for 2 h at room temperature with donkey anti-rabbit secondary antibody conjugated to Alexa Fluor 488, diluted 1:500 in TBST containing 1%. Sections were then washed, mounted onto glass slides, and coverslipped with vectashield mounting medium (Vector Labs, H-1000). Negative controls were processed by omission of the primary antibody.

### 2.6 Image acquisition and quantification

Connexin immunoreactivity was quantified as puncta density within defined regions of interest (ROI) in the adrenal medulla. Images were acquired using a Zeiss LSM 880 confocal microscope. Z-stack images were captured with a 63× objective using an Airy unit setting of 1, resulting in an optical section thickness of <1 µm. For each marker, all images included in the same analysis were acquired using identical laser power, detector gain, offset, pinhole, scan speed, and image size settings.

For puncta analysis, ROI of 25 µm^2^were placed within chromaffin cells that were clearly identifiable by a dark nucleus and well-defined cellular boundaries. Approximately 5–10 cells were analyzed per section, and adrenal sections were acquired from at least 5 animals per experimental group. Connexin-positive puncta were quantified within each region of interest using FIJI, after applying a constant threshold across all images from the same staining batch. Puncta density was expressed as the number of immunoreactive puncta per 25 µm^2^.

### 2.7 Statistical analysis

Data are presented as mean ± SEM. Connexin-positive puncta density was analyzed separately by sex and connexin using one-way ANOVA followed by Tukey’s multiple-comparison test. Plasma corticosterone concentrations were analyzed using one-way ANOVA followed by Fisher’s LSD post hoc test. Normality of residuals was assessed using D’Agostino–Pearson. Differences were considered statistically significant when p < 0.05. Statistical analyses were performed using GraphPad Prism version 11.

## 3. Results

### 3.1 Unpredictable mild stress (UMS) increased adrenal medullary Cx36 and Cx43 immunoreactivity and neonatal maternal separation amplified this response

We first examined whether neonatal maternal separation produced a persistent basal change in connexin immunoreactivity in the adult adrenal medulla. Under basal conditions, Cx36 and Cx43 immunoreactivity appeared comparable between animal-facility-reared control animals (AFR) and maternally separated animals (MS) in both males and females (Figure 1). In males, Cx36-positive puncta density was 20.29 ± 0.79 puncta/25 µm^2^in AFR basal animals and 19.23 ± 0.57 puncta/25 µm^2^in MS basal animals. In females, Cx36-positive puncta density was 21.65 ± 0.73 puncta/25 µm^2^in AFR basal animals and 21.18 ± 1.01 puncta/25 µm^2^in MS basal animals. Neither comparison was statistically significant. A similar pattern was observed for Cx43. In males, Cx43-positive puncta density was 13.43 ± 0.47 puncta/25 µm^2^in AFR basal animals and 14.00 ± 0.41 puncta/25 µm^2^in MS basal animals. In females, Cx43-positive puncta density was 16.89 ± 0.33 puncta/25 µm^2^in AFR basal animals and 16.50 ± 0.50 puncta/25 µm^2^in MS basal animals. These basal comparisons were also not statistically significant. Thus, neonatal maternal separation did not produce a constitutive increase in adult adrenal medullary Cx36 or Cx43 immunoreactivity.

**Figure 1.**
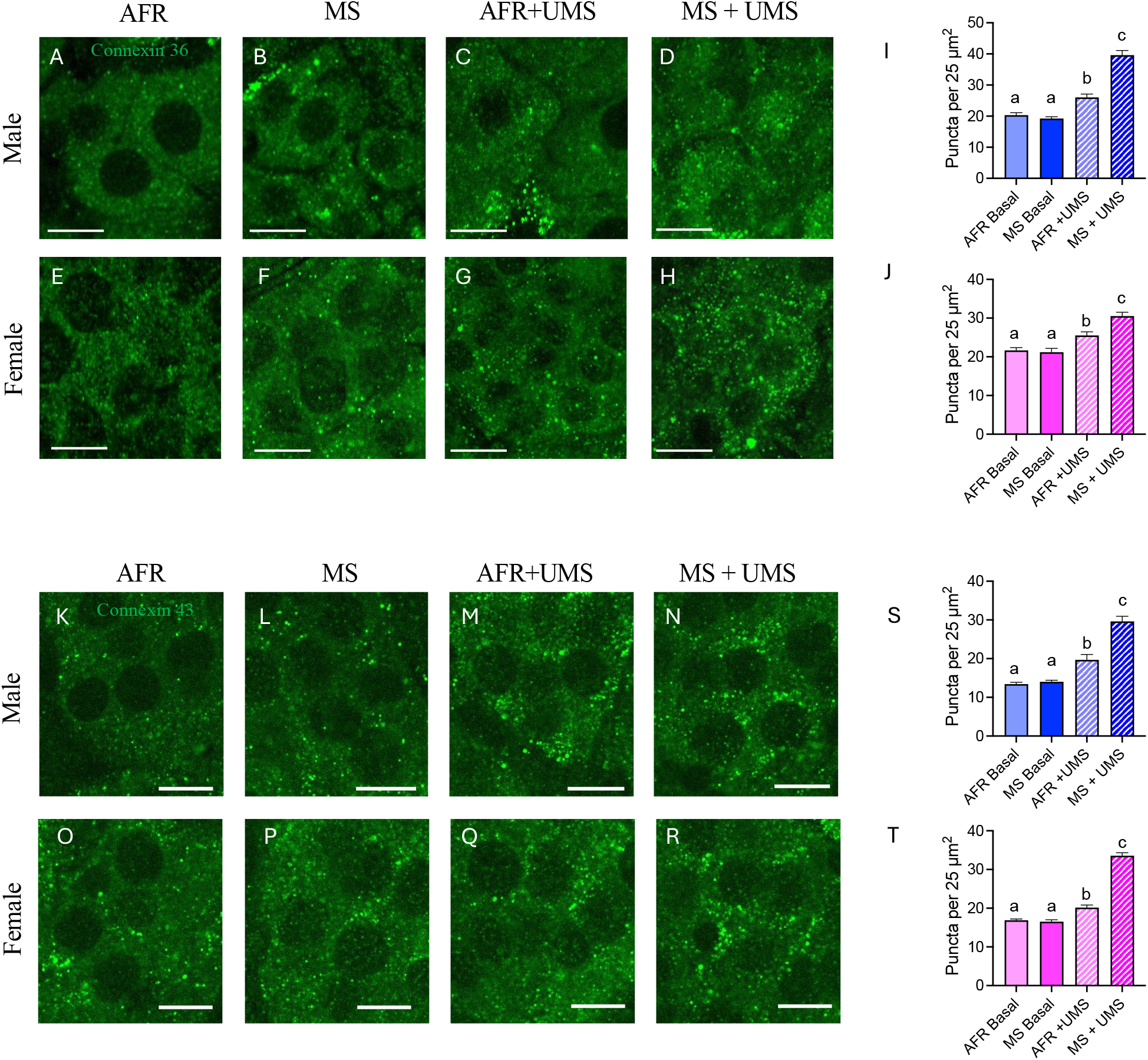
Neonatal maternal separation potentiates unpredictable mild stress-induced Cx36 and Cx43 immunoreactivity in the adult adrenal medulla. Representative confocal images and quantitative analysis of connexin 36 (Cx36) and connexin 43 (Cx43) immunoreactivity in the adrenal medulla of animal-facility-reared control rats (AFR) and maternally separated rats (MS), under basal conditions or after exposure to unpredictable mild stress (UMS). Panels A–H show Cx36 immunoreactivity. Bar graphs in panels I and J show quantification of Cx36 puncta per 25 µm^2^. Panels K–R show Cx43 immunoreactivity. For each connexin, the upper row corresponds to male rats and the lower row to female rats. Bar graphs in panels S and T show mean ± SEM of Cx43 puncta density expressed as puncta per 25 µm^2^. Different letters indicate statistically significant differences between groups (p < 0.05, one-way ANOVA followed by Tukey’s multiple-comparison test). Scale bar = 10 µm.

In contrast, adult UMS increased both Cx36 and Cx43 immunoreactivity in the adrenal medulla, and this response was consistently amplified in animals previously exposed to neonatal maternal separation. In males, Cx36 positive puncta density differed significantly among groups (one-way ANOVA, F(3,80) = 52.6, p < 0.001). AFR+UMS males showed higher Cx36 immunoreactivity than AFR basal males (26.00 ± 1.09 vs.

20.29 ± 0.79 puncta/25 µm^2^; Tukey post hoc test, p < 0.05). However, the largest increase was observed in MS+UMS males, which showed significantly higher Cx36 puncta density than AFR basal, MS basal, and AFR+UMS animals (39.60 ± 1.52 puncta/25 µm^2^; all p < 0.001).

Male Cx43 immunoreactivity showed the same general pattern. Cx43-positive puncta density differed significantly among groups (one-way ANOVA, F(3,78) = 34.1, p < 0.001). AFR+UMS males showed increased Cx43 immunoreactivity compared with AFR basal males (19.68 ± 1.40 vs. 13.43 ± 0.47 puncta/25 µm^2^; p < 0.05). MS+UMS males showed the highest Cx43 signal in this sex (29.60 ± 1.35 puncta/25 µm^2^), differing significantly from AFR basal, MS basal, and AFR+UMS groups (all p < 0.001).

In females, Cx36-positive puncta density also differed significantly among groups (one-way ANOVA, F(3,82) = 18.2, p < 0.001). AFR+UMS females showed higher Cx36 immunoreactivity than AFR basal females (25.51 ± 0.93 vs. 21.65 ± 0.73 puncta/25 µm^2^; p < 0.05). MS+UMS females showed the highest Cx36 signal in this sex (30.50 ± 1.01 puncta/25 µm^2^), differing significantly from AFR basal, MS basal, and AFR+UMS females.

Female Cx43-positive puncta density also differed significantly among groups (one-way ANOVA, F(3,77) = 139, p < 0.001). AFR+UMS females showed increased Cx43 immunoreactivity compared with AFR basal females (20.16 ± 0.65 vs. 16.89 ± 0.33 puncta/25 µm^2^; p < 0.01). MS+UMS females showed a marked increase in Cx43 immunoreactivity (33.54 ± 0.77 puncta/25 µm^2^), differing significantly from AFR basal, MS basal, and AFR+UMS groups.

Together, these results indicate that neonatal maternal separation did not produce a basal increase in adrenal medullary connexin immunoreactivity. Instead, adult UMS increased Cx36 and Cx43 puncta density, and this increase was consistently greater in animals with a history of neonatal maternal separation.

### 3.2 Maternal separation and unpredictable mild stress enhanced corticosterone responses to acute restraint

To determine whether stress-history-dependent connexin remodeling occurred in parallel with altered endocrine responsiveness, plasma corticosterone was measured under basal conditions and after 30 min of restraint stress, with or without prior exposure to the unpredictable mild stress protocol (Figure 2). Residuals from the corticosterone analysis passed D’Agostino–Pearson normality tests. Corticosterone concentrations differed significantly among experimental groups (one-way ANOVA, F(11,82) = 50.80, p < 0.0001), see Figure 2.

**Figure 2.**
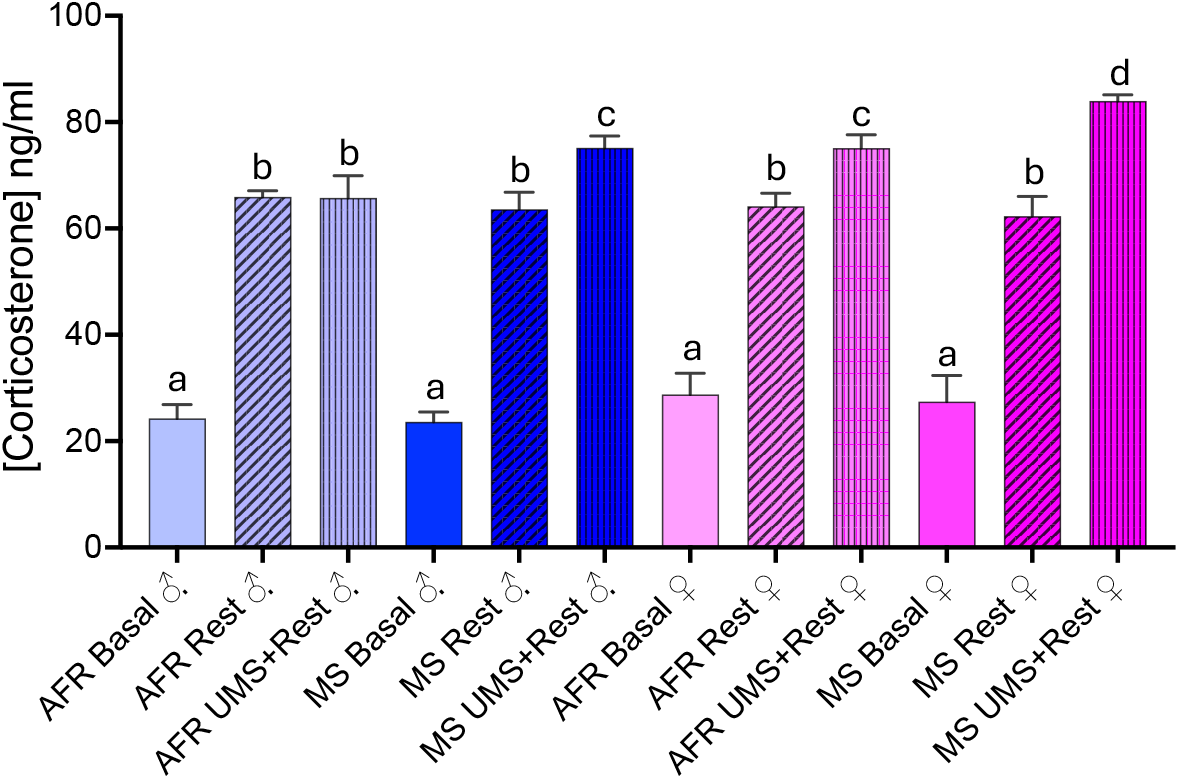
Neonatal maternal separation potentiates corticosterone responses after adult unpredictable mild stress. Plasma corticosterone concentrations were measured under basal conditions or after 30 min of restraint stress in adult animal-facility-reared control rats (AFR) and maternally separated rats (MS), with or without prior exposure to unpredictable mild stress (UMS). Maternal separation alone did not alter basal corticosterone concentrations or the corticosterone response to acute restraint. However, after UMS exposure, MS animals showed greater restraint-induced corticosterone responses than AFR controls, with the highest values observed in MS+UMS females. Data are shown as mean ± SEM. Different letters indicate statistically significant differences between groups (p < 0.05).

Basal corticosterone concentrations were low and comparable between AFR and MS animals in both sexes. In males, corticosterone was 24.27 ± 2.59 ng/mL in AFR basal animals and 23.61 ± 1.89 ng/mL in MS basal animals. In females, corticosterone was 28.80 ± 3.97 ng/mL in AFR basal animals and 27.46 ± 4.91 ng/mL in MS basal animals. These basal comparisons were not statistically significant, indicating that neonatal maternal separation alone did not produce a persistent elevation of basal corticosterone in adulthood.

Acute restraint increased corticosterone concentrations in both sexes. In males, corticosterone increased to 65.95 ± 1.15 ng/mL in AFR restraint animals and to 63.59 ± 3.22 ng/mL in MS restraint animals. These values did not differ significantly from each other. However, after prior UMS exposure, MS males showed higher corticosterone responses to restraint than AFR UMS+restraint males (75.16 ± 2.22 vs. 65.73 ± 4.19 ng/mL; p < 0.05). MS UMS+restraint males also showed higher corticosterone concentrations than MS restraint males without UMS (p < 0.01).

In females, restraint increased corticosterone to 64.18 ± 2.44 ng/mL in AFR animals and to 62.33 ± 3.70 ng/mL in MS animals, with no significant difference between these groups. Prior UMS further increased restraint-induced corticosterone, with AFR UMS+restraint females reaching 75.09 ± 2.54 ng/mL and MS UMS+restraint females reaching 83.98 ± 1.16 ng/mL. The MS UMS+restraint female group showed the highest corticosterone concentration of all groups and differed significantly from AFR UMS+restraint females, MS restraint females, and MS UMS+restraint males.

Overall, these findings indicate that neonatal maternal separation did not alter basal adult corticosterone or the corticosterone response to acute restraint alone. However, when followed by adult UMS exposure, prior maternal separation enhanced the corticosterone response to restraint. This endocrine phenotype paralleled the connexin findings, in which the combined history of early-life maternal separation and adult stress produced the strongest adrenal medullary remodeling and the highest corticosterone response.

## 4. Discussion

The present study shows that neonatal maternal separation primes adrenal medullary connexin remodeling in response to adult unpredictable mild stress. Neonatal maternal separation did not induced a constitutive increase in Cx36 or Cx43 puncta density in the adult adrenal medulla under basal conditions. However, the effect of early-life experience became evident after a later adult challenge: unpredictable mild stress increased adrenal medullary Cx36 and Cx43 immunoreactivity, and this response was consistently greater in animals previously exposed to maternal separation. This pattern suggests that maternal separation did not produced a permanently activated adrenal medullary phenotype, but rather modified the future capacity of chromaffin-cell networks to remodel in response to subsequent stress. In this sense, the data are consistent with a metaplastic model of developmental programming, in which early-life experience changes the threshold or magnitude of later plastic responses without necessarily producing evident basal alterations.

Gap junctional communication among chromaffin cells is well positioned to act as a local amplifier of adrenal medullary output. In adrenal slices, gap junctions mediate electrotonic communication between neighboring chromaffin cells, propagate cytosolic Ca^2+^increases, and allow stimulation of one chromaffin cell to recruit catecholamine release from coupled neighbors (Martin, Mathieu et al. 2001, Colomer, Olivos Ore et al. 2008). Subsequent work showed that stress can remodel this system. In cold-stressed rats, chromaffin cells exhibit increased dye coupling, stronger electrical coupling, action-potential transmission between coupled cells, increased Cx36/Cx43 protein expression, and enhanced catecholamine secretion that is sensitive to gap-junction blockade (Colomer et al., 2008). In vivo evidence further supports the physiological relevance of this mechanism: pharmacological blockade of adrenal gap junctional signaling reduces splanchnic nerve stimulation-evoked catecholamine release, with a larger effect after stress, and Cx36 deficiency impairs evoked secretion in mice (Desarmenien, Jourdan et al. 2013). The present findings extend this framework by suggesting that the magnitude of stress-induced connexin recruitment is influenced by early-life history.

The present findings support a model in which unpredictable mild stress (UMS) converts the adrenal medulla into a more strongly integrated chromaffin-cell network, characterized by increased Cx36 and Cx43 puncta density. Maternal separation amplifies this transition, indicating that early-life experience programs the future recruitment of gap-junctional communication during adult stress. In this framework, connexin remodeling is not merely a marker of adrenal activation, but a cellular mechanism through which the adrenal medulla scales its output according to prior stress history.

The current findings also fit with evidence that chromaffin-cell coupling is dynamically regulated by the state of splanchnic input. Cholinergic transmission and gap-junctional communication appear to be reciprocally regulated. Reduced or blocked cholinergic activity can increase gap-junctional coupling between chromaffin cells, suggesting tonic inhibitory control by cholinergic synaptic input (Martin et al., 2003). In addition, α9-containing nicotinic receptors participate in cholinergic synaptic transmission in the rat adrenal medulla and exert inhibitory control over dye coupling; these receptors are also remodeled by stress (Colomer et al., 2010). These studies support the idea that chromaffin-cell gap junctions are not passive anatomical features, but adaptable elements of adrenal stimulus–secretion coupling. PACAP provides a strong mechanistic link between adult stress exposure and the recruitment of chromaffin-cell network coupling. During elevated splanchnic activity, PACAP sustains catecholamine release through PLC/PKC-dependent facilitation of Ca^2+^entry (Kuri, Chan et al. 2009) and is required for both catecholamine secretion and transcriptional replenishment of the secretory phenotype during “stressed” high-frequency splanchnic stimulation (Stroth, Kuri et al. 2013). Importantly, enhanced electrical coupling and spread of excitation in mouse adrenal medullary slices (Hill, Lee et al. 2012), suggests that PACAP can act as an upstream modulator of the connexin-dependent coordination of chromaffin-cell activity (Zhang and Eiden 2019) . Within this framework, UMS may drive the adrenal medulla toward a PACAP-dominated, high-output state in which Cx36- and Cx43-dependent intercellular communication is increasingly recruited to synchronize chromaffin-cell activity. The larger connexin response observed in MS+UMS animals suggests that early-life stress primes this PACAP-connexin axis, allowing the adrenal medulla to engage a more integrated secretory network during adult stress.

A central mechanism linking maternal separation to later adrenal medullary plasticity may involve long-lasting changes in hypothalamic autonomic control. Previous work using the same maternal separation paradigm demonstrated downregulated programmed cell death in hypothalamus (Irles, Nava-Kopp et al. 2014) and a persistent remodeling of the hypothalamic vasopressin system, including increased AVP expression and enlargement of AVP-containing hypothalamic nuclei in adulthood (Zhang, Hernandez et al. 2012). The hypothalamic paraventricular nucleus (PVN) contains closely intermingled neuroendocrine and presympathetic neuronal populations that can influence autonomic outflow. Dendritically released vasopressin can mediate crosstalk between magnocellular neurosecretory neurons and preautonomic PVN networks, providing a route through which vasopressinergic activity may enhance sympathetic recruitment (Son, Filosa et al. 2013). Consistent with this anatomical framework, our representative AVP/V1a immunohistochemical observations show V1a-positive elements within the PVN in close proximity to AVP-positive neuronal profiles and processes.

Sex may shape the adrenal consequences of combined early-life and adult stress exposure. In the present dataset, both males and females showed increased Cx36 and Cx43 immunoreactivity after UMS, and both sexes showed larger responses after the combined MS+UMS condition. The corticosterone data showed the clearest sex-associated pattern, with the highest endocrine response observed in MS+UMS females. These findings suggest that females may be particularly sensitive to the combined effect of neonatal maternal separation and adult stress exposure (Zetter, Hernandez et al. 2021). Future studies using factorial analyses designed to test sex as an independent variable, together with estrous-cycle monitoring or gonadal hormone manipulation, will be needed to define the mechanisms underlying these sex-associated differences.

Future studies should determine how the connexin remodeling described here modifies adrenal medullary function at the network level. The increase in Cx36 and Cx43 puncta density after UMS, particularly in MS+UMS animals, suggests enhanced recruitment of gap-junctional communication within chromaffin-cell clusters. Combining connexin imaging with dye coupling, electrophysiology, Ca^2+^imaging, catecholamine secretion assays, and selective manipulation of PACAP signaling will be important to define how this structural remodeling translates into coordinated secretory output. In addition, future studies should determine whether PVN vasopressinergic signaling, splanchnic nerve activity, and adrenal PACAP release form a functional axis linking early-life stress to enhanced chromaffin-cell network recruitment during adult challenge.

In conclusion, neonatal maternal separation programs the adult adrenal medulla to recruit connexin-dependent chromaffin-cell network remodeling more strongly during later unpredictable mild stress. This programming is not evident as tonic adrenal activation under basal conditions, but is unmasked by adult challenge, consistent with a metaplastic form of developmental stress adaptation. Together with the enhanced corticosterone response observed in MS+UMS animals, these findings support a model in which early-life stress biases hypothalamic–sympathoadrenal circuits toward amplified adrenal recruitment. Chromaffin-cell connexin remodeling may therefore constitute a previously underappreciated cellular mechanism by which early-life experience shapes the intensity and coordination of adult neuroendocrine stress responses.

## Acknowledgement

This work was supported by the Mexican Secretaría de Ciencia, Humanidades, Tecnología e Innovación (Secihti) through grants CF-2023-G-243 and CIORGANISMOS-2025-92 to LZ, DGAPA-PAPIIT-UNAM IT200125 to VSH. We thank Dr. Lee E. Eiden (National Institute of Mental Health, NIH, USA) for generously providing connexin antibodies used in this study and for valuable scientific discussions. We are grateful to María José Gomora and Lorena Mejía-Barretto for excellent technical assistance.

